# Predictive coding is a consequence of energy efficiency in recurrent neural networks

**DOI:** 10.1101/2021.02.16.430904

**Authors:** Abdullahi Ali, Nasir Ahmad, Elgar de Groot, Marcel A. J. van Gerven, Tim C. Kietzmann

**Affiliations:** Donders Institute for Brain, Cognition and Behaviour, Radboud University, Nijmegen, the Netherlands; Department of Experimental Psychology, Utrecht University, Utrecht, the Netherlands

## Abstract

Predictive coding represents a promising framework for understanding brain function. It postulates that the brain continuously inhibits predictable sensory input, ensuring a preferential processing of surprising elements. A central aspect of this view is its hierarchical connectivity, involving recurrent message passing between excitatory bottom-up signals and inhibitory top-down feedback. Here we use computational modelling to demonstrate that such architectural hard-wiring is not necessary. Rather, predictive coding is shown to emerge as a consequence of energy efficiency. When training recurrent neural networks to minimise their energy consumption while operating in predictive environments, the networks self-organise into prediction and error units with appropriate inhibitory and excitatory interconnections, and learn to inhibit predictable sensory input. Moving beyond the view of purely top-down driven predictions, we furthermore demonstrate, via virtual lesioning experiments, that networks perform predictions on two timescales: fast lateral predictions among sensory units, and slower prediction cycles that integrate evidence over time.

## 1 Introduction

In computational neuroscience and beyond, predictive coding has emerged as a prominent theory for how the brain encodes and processes sensory information (Srinivasan et al., 1982; Mumford, 1992; Friston, 2005; Clark, 2013). It postulates that higher-level brain areas continuously keep track of the causes of lower-level sensory input, and actively inhibit incoming sensory signals that match expectation. Over time, the brain’s higher-level representations are shaped to yield increasingly accurate predictions that, in turn, minimise the surprise that the system encounters in its inputs. That is, the brain creates an increasingly accurate model of the external world and focuses on processing unexpected sensory events that yield the highest information gain.

Adding to the increasing, albeit often indirect, experimental evidence for predictive coding in the brain (Alink et al., 2010; Näätänen et al., 2001; Summerfield et al., 2011, 2008; Squires et al., 1975; Hupé et al., 1998; Murray et al., 2002; Rao et al., 2016; Kok et al., 2012; Ekman et al., 2017; De Lange et al., 2018; Dijkstra et al., 2020; Schwiedrzik and Freiwald, 2017), computational modelling has investigated explicit implementations of predictive coding, indicating that they can reproduce experimentally observed neural phenomena (Rao and Ballard, 1999; Lee and Mumford, 2003; Friston, 2005, 2010). For example, Rao and Ballard (1999) demonstrated that a hierarchical network with top-down inhibitory feedback can explain extra-classical field effects in V1. In addition, deep neural network architectures wired to implement predictive coding were shown to work at scale in real-world tasks (Chalasani and Principe, 2013; Lotter et al., 2016; Villegas et al., 2017; Lotter et al., 2020), adding further support for the computational benefits of recurrent message passing (Spoerer et al., 2020; Kietzmann et al., 2019; Kar et al., 2019; Linsley et al., 2020; Spoerer et al., 2017; van Bergen and Kriegeskorte, 2020). The common rationale of predictive coding focused modelling work is to test for its computational and representational effects by hard-wiring the model circuitry to mirror hierarchical and inhibitory connectivity between units that drive predictions, and units that signal deviations from said predictions (also termed sensory, or error units).

Contrasting this approach, we here ask a different question: can computational mechanisms of predictive coding naturally emerge from other, perhaps simpler, computational principles? In search for such principles, we take inspiration from the organism’s limited energy budget. Expanding on previous work on efficient coding and sparse coding (Barlow and Rosenblith, 1961; Bell and Sejnowski, 1995; Olshausen and Field, 1996; Bialek et al., 2006; Berkes and Wiskott, 2005; Chalk et al., 2018; Eckmann et al., 2020), we subject recurrent neural networks (rate-based RNNs) to predictable sequences of visual input and optimise their synaptic weights to minimise what constitutes the largest sources of energy consumption in biological systems: action potential generation and synaptic transmission (Sengupta et al., 2010). We then test the resulting networks for phenomena typically associated with predictive coding. We compare the energy budget of the networks to theoretically-determined lower bounds, search for inhibitory connectivity profiles that mirror sensory predictions, and explore whether the networks automatically separate their neural resources into prediction and error units.

## 2 Results

To better understand whether, and under which conditions, predictive coding may emerge in energy-constrained neural networks, we first trained a set of RNNs on predictable streams of visual input. For this, we created sequences of handwritten digits taken from the MNIST data set (LeCun et al., 1998) iterating through 10 digits in numerical (category) order. Starting from a random digit, sequences wrapped around from nine to zero. The visual input to each network unit was stimulus-dependent, with each unit receiving an input which mirrored its corresponding pixel intensity (i.e. 784 network units, corresponding to 784 input pixels from MNIST). These visual inputs were not weighted with modifiable parameters, preventing the models from learning to ignore the input as a shortcut to a solution for energy efficiency. Instead, we hypothesised that the models would learn to use their recurrent connectivity to counteract the input drive where possible, and thereby to minimise their energy consumption. Energy costs were defined to arise from synaptic transmission and action potentials, thereby mirroring the two most dominant sources of energy consumption in neural communication (Sengupta et al., 2010). Figure 1 shows an overview of the experimental pipeline. Further details are provided in sections 4.1 and 4.2.

**Figure 1:**
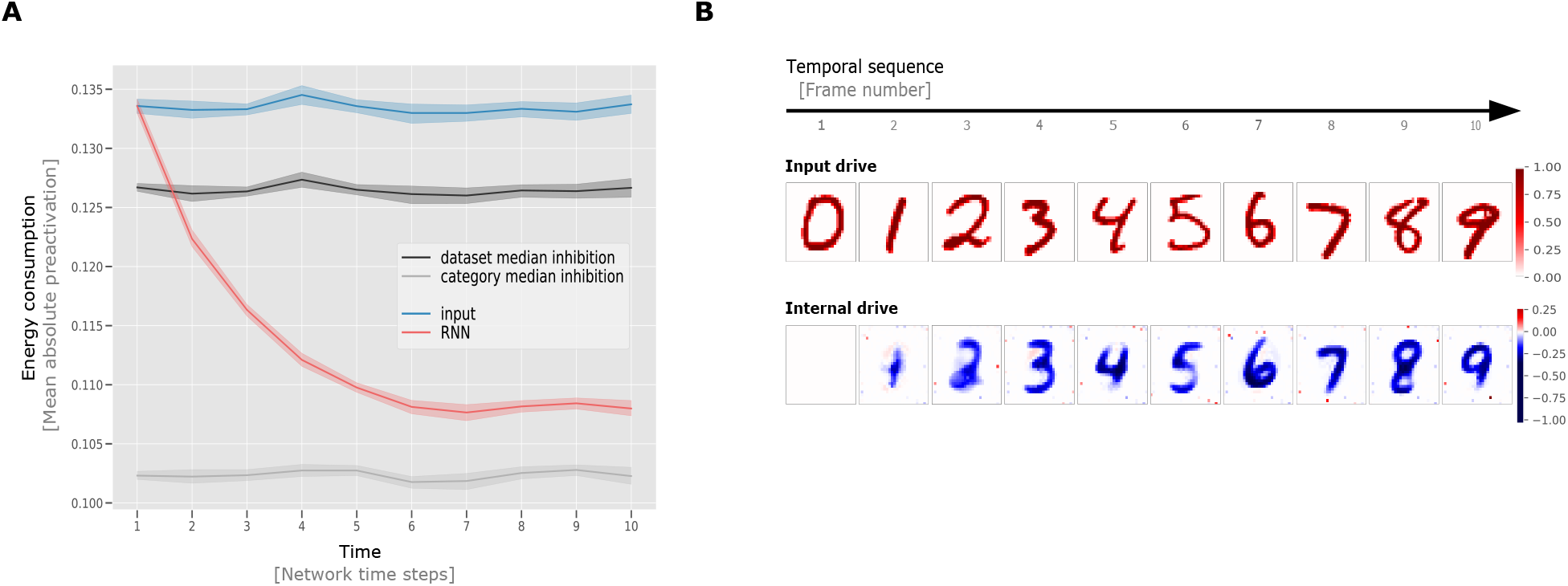
Overview of the model architecture and input pipeline. The upper part of the figure shows an example input sequence fed to a recurrent network. In each run, 10 randomly sampled MNIST digits are presented in ascending order to the network (with wraparound after digit 9). Each input pixel drives exactly one unit of the RNN. The preactivation of each RNN unit is computed from the recurrent input, provided by other networks units, as well as the input drive determined by the intensity of the pixel that it is connected to. The output of the network units is determined by taking the ReLU of the preactivations.

### 2.1 Preactivation as a proxy for the energy demands of the brain

Sparse coding is often implemented as an L1 objective on the unit outputs, i.e. networks are optimised to reduce the overall unit activity post non-linearity by having as few units active as possible. However, given that synaptic transmission contributes considerable energy cost alongside action potentials, a different learning objective is required to approximate total energy consumption; for example, a minimisation of unit outputs alone could be implemented via unreasonably strong inhibitory connections, which themselves might increase the overall energy budget. To solve this issue, we propose to instead train energy-efficient RNNs by minimising absolute (L1) unit preactivation. This has two desired properties. First, it drives unit output towards zero, mirroring a minimisation of unit firing rates (dependent naturally upon the form of activation function used). Second, we show that this objective also leads to minimal synaptic weight magnitudes in cases of noise in the system, thereby mirroring an overall reduction in synaptic transmission (see Appendix B for a theoretical derivation of why this is the case). We consider this link between a biologically realistic notion of energy efficiency and synaptic weight magnitude an important first result of this paper.

Optimising for minimal preactivation, we trained ten network instances with different random weight initialisations (Mehrer et al., 2020). The resulting network activity was compared to three theoretical scenarios. First, we considered a scenario in which the lateral weights are removed from the network, and the RNNs are therefore completely driven by external input. This measures the “overall input drive”, i.e. the energy that flows into the network from sensory sources alone. Second, we considered a scenario in which the networks counteract the input drive based on the global data set statistic, without sequence-specific categorical inhibitions. For this, we estimated the theoretical network activity if RNNs learned to predict the pixel-wise median value of the whole training data, irrespective of temporal order. Third, we considered a best case scenario in which the RNNs learn to predict and inhibit the pixel-wise median value of the particular digit category being presented. This theoretical model has perfect knowledge of the current position in the input sequence and can therefore inhibit the matching category pixel-wise median. Given the current setup, this measure represents a lower bound on our loss function, namely the summed absolute network preactivation (energy). A derivation for these bounds on performance is given in Section 4.3. All empirical network data shown is derived from a test data set that the networks were not trained on.

As can be seen in Figure 2A, which depicts the theoretical bounds together with the average preactivation across ten trained RNN instances, the trained networks approach the lower energy bound with increasing network time (i.e. increasing exposure to the test sequence). Whereas the RNNs cannot predict well at the first time step due to the random sequence onset, the average unit preactivation drops below the pixel-wise median of the training data after only one timestep and subsequently converge towards the category median, i.e. the lower bound. This indicates that the RNNs integrate evidence over time, improving their inhibitory predictions with increasing sensory evidence for the current input state.

**Figure 2:**
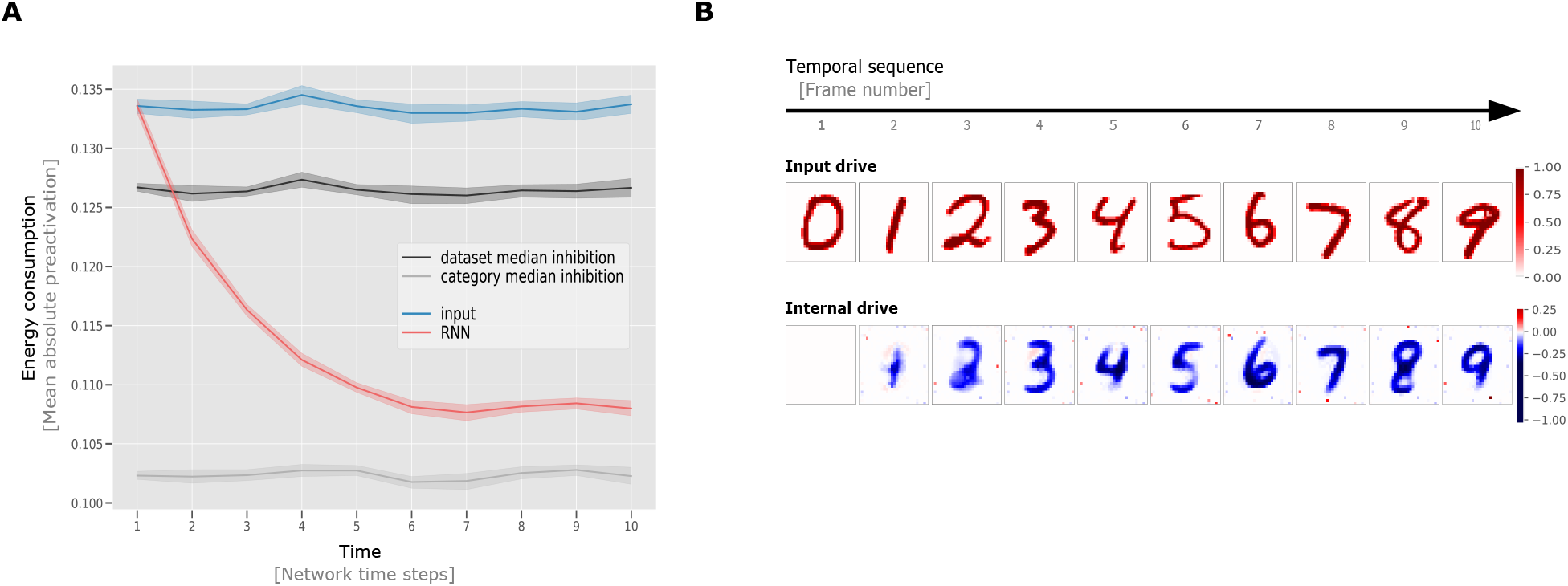
Network performance and behaviour. **A:** Evolution of the ‘energy consumption’ (mean absolute preactivation) of the RNNs when presented with test data, together with three theoretical scenarios. *input*: stimulation entering the networks via sensory input, *median*: network preactivation if the networks predicted and inhibited the (pixel-wise) median of the training data set, *category median*: network preactivation if the networks had perfect knowledge of the current state of the sequence and thereby inhibit the (pixel-wise) median value of the upcoming digit category (lower energy bound). *RNN*: actual network preactivation. Mean preactivations shown with 95% confidence intervals, bootstrapped across 10 network instances with replacement. As can be observed, the (RNN) network preactivations achieve lower energy with longer exposure to stimuli, suggesting an integration of evidence over time. **B:** Network activity for an example sequence. The RNN units are organised according to the locations of their input pixels. Row 1 shows the input drive and row 2 shows the internal drive. Excitatory signals are depicted in red, while inhibitory signals are depicted in blue. The darker the colour the stronger the signal. The sequence images shown were not used for training. The observable improvements in inhibitory network drive indicate that the RNN integrates sensory evidence over time, leading to better internal knowledge of the state of the world, and more targeted inhibitory predictions.

### 2.2 RNNs trained for energy efficiency learn to predict and inhibit upcoming sensory signals

Having established that the trained RNNs successfully reduce their preactivation over time, we further confirmed a targeted inhibition of predicted sensory input, a hallmark of predictive coding. This was accomplished by separately visualising the input drive, i.e. the unit activation driven by sensory input, and the network drive, i.e. the unit activation due to network-internal connectivity alone. Figure 2B depicts an example sequence using test data (top row: input drive, bottom row: (recurrent) internal drive, see Section 4.2 for further details on these measures). Two observations are of importance in this example. First, the network clearly predicts and inhibits the upcoming digit category. Second, as time progresses the network is able to integrate information, leading to better knowledge of the current sequence position, and more targeted inhibitions (e.g. the inhibition of later digits appears more strong/targeted than earlier predictions that appear weaker/smoother). Better predictions, in turn, result in progressively lower network activity, as previously quantified. Please note that this pattern of results is in line with hierarchical predictive coding architectures in which feedback from higher order areas is subtracted from sensory evidence of lower areas. However, neither a hierarchical organisation nor activity subtraction was imposed in our RNNs. Rather, all effects emerge due to optimisation of the networks for energy efficiency.

### 2.3 Separate error units and prediction units as an emergent property of RNN self-organisation

Our previous results demonstrate that RNNs, when optimised to reduce their energy consumption (action potentials and synaptic transmission), exhibit key phenomena associated with predictive coding. Predictive coding emerges, without architectural hard-wiring, as a result of energy efficiency in predictable sensory environments. A second key component postulated by the predictive coding framework is the existence of distinct neural populations that either provide sensory predictions, or which carry the deviations from such predictions, i.e. prediction error units (Bastos et al., 2012). Given the non-hierarchical, non-constrained nature of the current setup, we next asked whether a similar separation could also be observed in our energy efficient networks.

If the models had indeed developed separate populations of error and prediction units, they would be expected to have differences in their median preactivations in the latter time points of the sequence (i.e. the time when network dynamics should be most stable). Given that the networks are trained to reduce preactivation of their units, any units with consistent nonzero median preactivation are in line with the interpretation as having a functional role in driving down preactivation of other units. Following this rationale, we identified candidates for prediction units by calculating the median preactivation of each unit across multiple instances of each digit category, during the presentation of the final element of the sequences. Units were labelled to be potentially predictive, if for any one class, the 99% confidence interval (*CI*) around the median preactivation did not include zero. That is, they showed a consistently larger than zero activity profile, which would have been inhibited if it did not serve a functional role in the overall objective of energy efficiency. See 4.4 for details on the exact procedure for categorising units and calculation of CI’s for the median preactivation and median absolute deviation (MAD) around the median preactivation. Results of this analysis for an example digit category (digit 9) is shown in Figure 3A. As can be seen from the example scatter plot, contrasting the units’ median preactivation and median absolute deviation (MAD) around that median, two populations emerge. Scatter plots of the other classes are provided in Appendix A.

**Figure 3:**
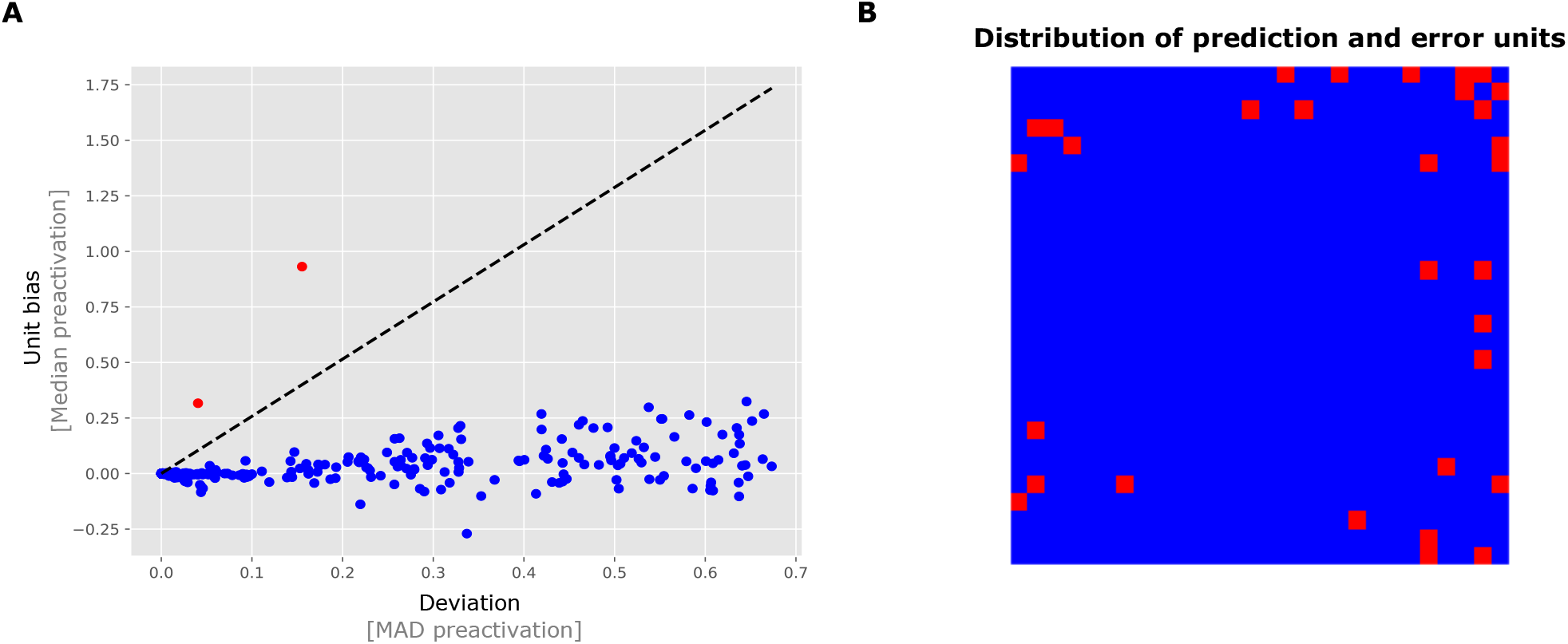
Trained RNNs exhibit distinct populations of units: units with approximately zero median preactivation and units with a ‘bias’ in their median preactivation larger than zero. **A:** a scatter plot for digit 9 of an example network indicates a separation between units based on their median preactivation. Blue dots indicate units for which their 99% CI around the median includes zero. Red dots indicate the units for which their 99% CI around median did not cross zero. The dashed line depicts the decision boundary that separates the two populations (only shown in the positive direction of interest). **B:** topographical distribution of populations in MNIST space. Units that had nonzero median preactivation for at least one class were stained red, the remaining units were stained blue. The postulated prediction units (red squares) predominantly occur in the network/stimulus periphery.

Of particular note, when all potentially predictive units are collected they occupy a different topographic part of the RNN (see Figure 3B, which shows the general distribution of postulated prediction- and sensory units across all ten digit categories). Units with consistent positive activation (i.e. the postulated prediction units) reside in locations that are typically not strongly driven by sensory input, see the structure of the MNIST data set. Such secondary use of neural resources that would otherwise remain unused is in line with neuroscientific evidence for cortical plasticity and reorganization (Kaas, 2002; Merabet and Pascual-Leone, 2010).

Please note that until now, we have not established a causal effect or functional role of the postulated prediction units in the network dynamics. This will be addressed in depth in the next section.

### 2.4 Prediction units integrate evidence over time for more targeted inhibition

To explicitly test for their functional role in the RNN dynamics, we performed virtual lesion experiments on the candidate prediction units, as identified in the previous section. As shown in Figure 4B (top row), an un-lesioned RNN is capable of more fine-grained predictions with increasing sequence lengths leading up to the sensory input. In stark contrast to this, a network with lesioned prediction units (Figure 4B, bottom row) remains fixed at the initial and immediate prediction and misses the ability to integrate evidence over time (see Supplemental Figures SA1 and SA2 for further lesion experiments targeting other digit categories). Next, we quantified the network-wide effects of lesioning prediction units (Figure 4A). Following an initial reduction in energy consumption (investigated in depth in the next section), the lesioned network fails to reduce its activity over time (yellow line). As a control experiment, we lesioned the same number of units from outside the population of identified prediction units, rather than prediction units (green line). This had minimal effect on the network energy consumption, verifying the special role of prediction units in overall network dynamics.

**Figure 4:**
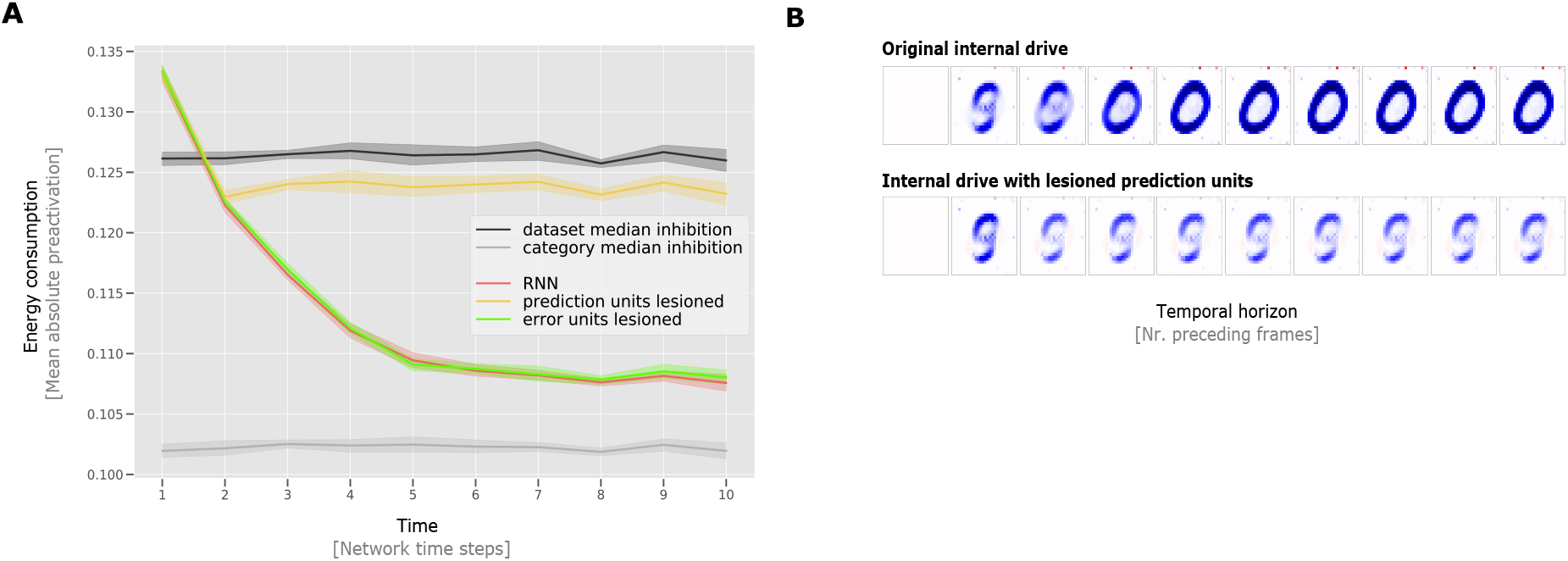
Lesion studies reveal the causal role of prediction units. **A:** mean preactivations of normal RNNs (red line), prediction unit lesioned RNNs (yellow line), and error unit lesioned RNNs (green line). In contrast to regular RNNs, the prediction unit lesioned RNNs do not reduce their preactivation throughout the sequence, indicating that they are not capable of integrating evidence over time. Control experiments demonstrate that lesioning error units instead of prediction units has little effect on network dynamics (green line). From this, we can conclude that the prediction units have a unique functional role in prediction. Mean preactivations shown with 95% confidence intervals, bootstrapped across 10 network instances with replacement. **B:** internal drive (*p*_*t*_) in the RNN for the ‘0’ digit category as a result of varying temporal history prior to the stimulus presentation. Each square shows the internal drive of the network following various sequence length prior to the stimulus of interest, e.g. the sixth image displays the internal drive after the sequence ‘5-6-7-8-9’ was presented to the network. The first row shows the predictions of a normal RNN, while the second row shows the result of lesioning RNN prediction units. As can be seen, the internal drive of the lesioned network does not develop better predictions with longer sequence lengths.

### 2.5 Two distinct inhibitory mechanisms work at different timescales

As can be seen in Figure 2, the energy consumption of the network drops from the first to the second timestep by means of an imprecise, yet category-specific inhibition. This network behaviour is of interest, as potential prediction loops, from input to prediction units and back, require two timesteps to come into effect (one to activate prediction units and one for these prediction units to provide feedback). This suggests that the observed predictions in timestep 2 are rather driven by more immediate lateral connections among error units - to our knowledge a mode of predictive coding not previously considered. In summary, we observe two different modes of predictive inhibitions: one operating at a faster timescale among error units, and one on a slower timescale involving prediction units with the potential to integrate evidence over time.

### 2.6 Replication of all main results: CIFAR10

Having established that predictive coding can emerge as a result of training RNNs for energy efficiency, we tested the generality of our results by testing on a separate data set of more realistic image statistics (e.g. full image coverage, compared to MNIST in which information is predominantly present in central pixels). In particular, we performed experiments as before, but used sequences of CIFAR10 categories with colour instead of MNIST (see figure 5). Replicating our previous results, we again observe the emergence of two distinct populations of units, with lesions to prediction-units exhibiting strong effects on the predictive performance and energy consumption of the network (figure 5 panel A). One notable difference is that we observe far fewer prediction units in CIFAR10 than in MNIST despite the networks being bigger (784 units vs. 3072 units). This is likely due to the fact that in the CIFAR10 data set all units are strongly driven by sensory input, which increases the complexity of the optimisation problem and prevents the network from developing class-specific prediction units. Nevertheless, the networks are still able too minimise preactivation and reproduce the lesioning results seen in the MNIST data set (Figure 5C). See section 4.1 for specific details on the CIFAR10 data set.

**Figure 5:**
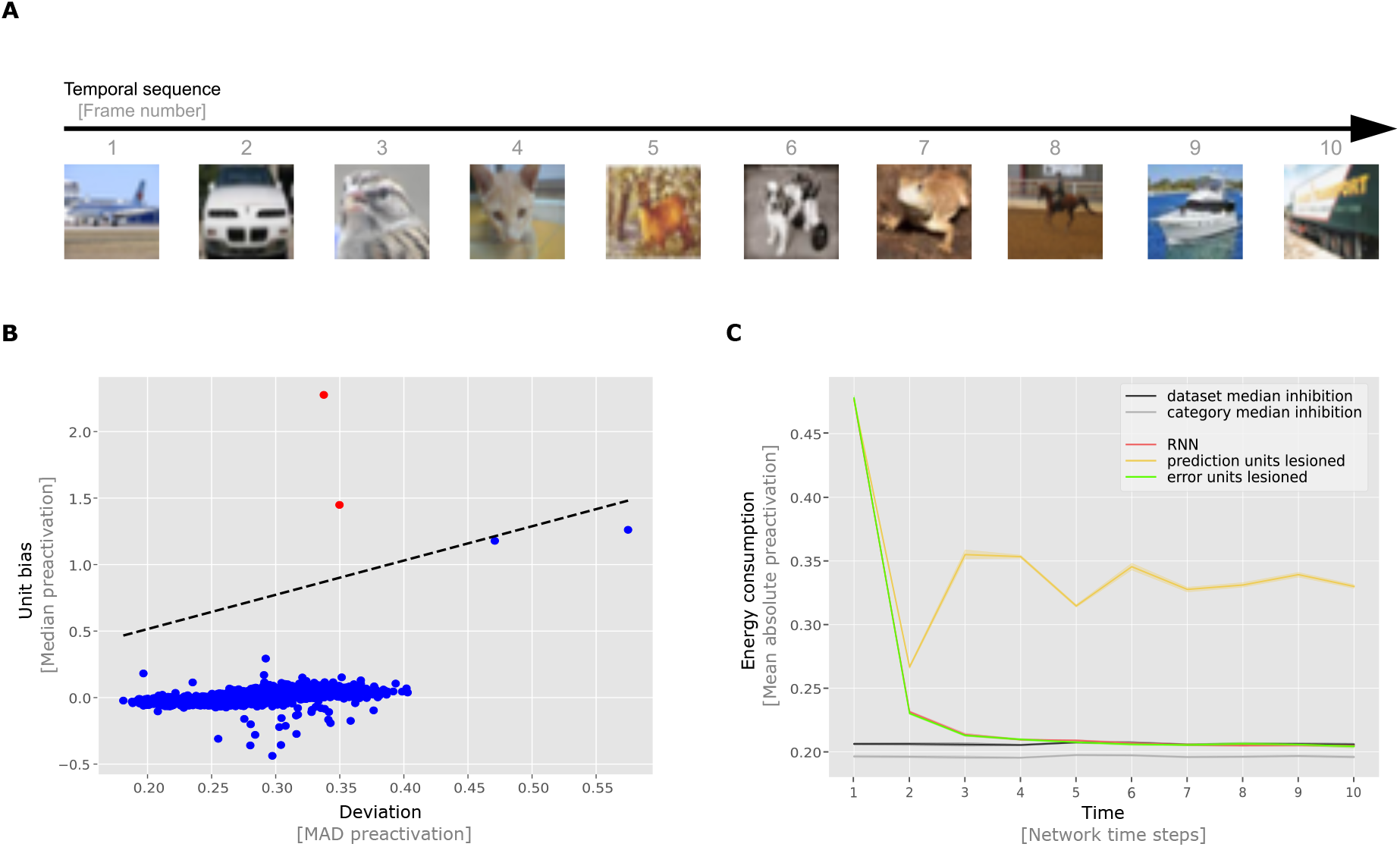
Results generalise to CIFAR10. **A:** example input sequence fed to the network. In each run 10 randomly sampled CIFAR10 images are presented in ascending order to the network (with wrap around after image class 9). **B:** scatter plot for the truck class of an example network indicates a separation between units based on their median preactivation. Blue dots indicate units for which their 99% CI around the median crossed zero. Red dots indicate the units for which their 99% CI around median did not cross zero. **C:** mean preactivations of normal RNN and lesioned RNNs. In contrast to normal RNNs, the lesioned RNNs do not reduce their preactivation over time, indicating that they are not capable of integrating evidence over time. Green line indicates control where error units are lesioned. Since lesioning the error units does not significantly increase network preactivation we can conclude that the prediction units have a unique functional role in prediction. Mean preactivations shown with 95% confidence intervals, bootstrapped across 10 network instances with replacement.

## 3 Discussion

Here we demonstrate that recurrent neural network models, trained to minimise biologically realistic energy consumption (action potentials and synaptic transmissions), spontaneously develop hallmarks of predictive coding without the need to hard-wire predictive coding principles into the network architecture. This opens up the possibility that predictive coding can be understood as a natural consequence of recurrent networks operating under a limited energy budget in predictive environments. In addition to category-specific inhibition and increasingly sharpened predictions with time, we observed that the RNNs self-organise into two distinct populations in line with error units and prediction units. Beyond these processes, we observe two distinct mechanisms of prediction, a faster inhibitory loop among error units, and a slower predictive loop that enables RNNs to integrate information over longer timescales and thereby to generate increasingly fine-grained predictions. This observation can be interpreted as a rudimentary form of hierarchical self-organisation in which the predictive units can be viewed as a higher-order cortical population operating on longer timescales and the error units as a lower-order cortical population operating on shorter timescales. This interpretation is consistent with hierarchical predictive coding architectures such as Rao and Ballard (1999) or Lotter et al. (2016) as well as the organisation of the brain in terms of a hierarchy of timescales (Kiebel et al., 2008; Baldassano et al., 2017). The phenomenon of specialisation in the circuit has also been demonstrated to be a useful notion by Zeldenrust et al. (2021), who analytically derived a class of spiking recurrent predictive coding networks with neural heterogeneity and has shown that these networks can better explain away prediction errors than homogeneous networks. This may indicate that neural heterogeneity and specialisation is an integral part of neural systems that have to be efficient in predictive environments.

Our loss function is computed from the network preactivation, and thereby fully depends on both the input drive and the recurrent drive. This implies an optimal solution in which the recurrent drive corresponds to the negative input drive. The network could therefore be seen as optimising for the negative target, and therefore within this context there is little to no difference between energy minimisation and prediction of network input. While it is true that minimising the preactivation is equivalent to predicting the (negative) input drive, this does not necessarily imply a predictive coding framework. In particular, predictive coding as a framework consists of functionally distinct population of units interacting in a hierarchical fashion Bastos et al. (2012). Thus, the loss function minimised, despite its functional relationship to prediction, does not build in these assumptions of structure. This allows us to conclude that our loss function, though intimately linked to prediction, does not assume the predictive coding framework and this is instead an emergent property This works builds upon a number of studies investigating the relationship between predictive systems, environments, and their impacts on efficiency. From a thermodynamics perspective, Still et al. (2012) demonstrated that a system with memory exposed to a stochastic signal must be predictive in order to operate at maximal energy efficiency. Candadai and Izquierdo (2020) showed, information-theoretically, that predictable environments produce neural networks which exhibit predictive information. On a neural level, it was demonstrated that tightly balanced excitation-inhibition can be understood as neural networks efficiently coding information, with membrane potentials of neurons interpretable as a prediction error of a global signal (Boerlin et al., 2013; Denève and Machens, 2016; Denève et al., 2017; Brendel et al., 2020). Masumori et al. (2019) demonstrate that a spiking neural network, solely trained based upon spike-timings, learns to predict temporal sequences, proposing that predictive coding arises in order to avoid stimulation of biological networks. These studies investigate prediction and efficiency at many scales and across systems. This current work extends this body of evidence by demonstrating that the framework for predictive coding can emerge in recurrent networks which are trained for a simple consideration of energy efficiency, reproducing functional components of predictive coding in a biologically-inspired network setup. It should be noted that the current work, due to its reliance on smaller-scale data sets (MNIST and CIFAR10) represents a proof of concept. The reliable findings across two separate data sets and the aforementioned findings on the link between prediction and energy efficiency indicates, however, that the observed results may be more general. Along this line of research, future work will explore larger-scale recurrent systems and their interplay with more natural image statistics. In terms of model details, future work could also explore energy efficiency in spiking network models where membrane voltages and spike-times must be considered, as the current set of results relies on a rate-coded neuron model. In addition, how a predictive system should optimally process differing spatial and temporal scales of dynamics and deals with unpredictable but information-less inputs (e.g. chaotic inputs and random noise) are key areas for future consideration.

In summary, our current set of findings suggests that predictive coding principles do not necessarily have to be hard-wired into the biological substrate but can be viewed as an emergent property of a simple recurrent system that minimises its energy consumption in a predictable environment. Notably, we have shown here that minimising unit preactivations, under mild assumptions of noise in the system, implies minimising both unit activity and synaptic transmission. This observation opens further interesting avenues of research into efficient coding and neural network modelling.

## 4 Materials and methods

### 4.1 Data generation

#### 4.1.1 MNIST

The input data are sequences of images drawn from the MNIST database of handwritten digits (LeCun et al., 1998). Images are of size 28 × 28 with pixel intensities in the range [0, 1]. There are 60,000 images in the training set and 10,000 images in the test set. Each set of images can be divided into 10 categories; one for each digit. The frequency of each digit category in the data set varies slightly (frequencies lie within 9% - 11% of the total data set (70,000 samples)). Sequences are generated by choosing random digits as starting points. Digits from numerically subsequent categories are then randomly sampled and appended to the sequence until the desired sequence length is reached. The categories are wrapped-around such that after category ‘nine’ the sequence continues from category ‘zero’. The sequence length can be chosen in advance, but all sequences are constrained to have the same length (in our simulations we take a sequence length of 10). The sequences are organised into batches. The images are drawn without replacement and the process is stopped when there are no image samples available for the next digit. Incomplete batches, in which there are no remaining image samples to complete a sequence, are discarded.

#### 4.1.2 CIFAR10

The input data are sequences of images drawn from CIFAR10, a labelled subset of the tiny image database (Krizhevsky and Hinton, 2009). CIFAR10 consists of 10 classes. These classes are: airplanes, cars, birds, cats, deer, dogs, frogs, horses, ships, and trucks. Images are coloured and of size 32 × 32 × 3 with pixel intensities in the range [0, 1]. There are 50,000 images in the training set and 10,000 images in the test set. Each set of images can be divided into 10 categories; one for each class. The frequency of each category in the data set is exactly 6000. Sequences are generated by choosing random images as starting points. images from numerically subsequent categories are then randomly sampled and appended to the sequence until the desired sequence length is reached. The categories are wrapped-around such that after category ‘nine’ the sequence continues from category ‘zero’. The sequence length can be chosen in advance, but all sequences are constrained to have the same length (in our simulations we take a sequence length of 10). The sequences are organised into batches. The images are drawn without replacement and the process is stopped when there are no image samples available for the next class. Incomplete batches, in which there are no remaining image samples to complete a sequence, are discarded.

### 4.2 RNN architecture and training procedure

We created a fully connected recurrent neural network (RNN) consisting of *N* = 784 units for MNIST and *N* = 3072 units for CIFAR10. Each unit is driven by exactly one input pixel, which means that the number of units exactly matches the number of pixels in the image. The equations that determine the RNN dynamics are:

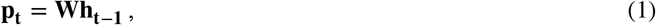

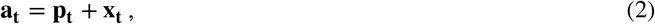

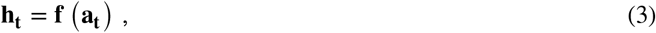

where **x**_**t**_ ∈ ℝ^*N*^ denotes the input drive, **a**_**t**_ ∈ ℝ^*N*^ denotes the *preactivation*, **W** ∈ ℝ^*N*×*N*^ denotes the recurrent weight matrix of the RNN (the only learnable parameters), **p**_**t**_ ∈ ℝ^*N*^ denotes the recurrent feedback, and **h**_**t**_ ∈ ℝ^*N*^ the unit outputs (in our study **f** are ReLU non-linearities). The subscripts, *t* and *t* − 1, refer to the discrete (integer) timestep of the system since all of these variables (aside from the weights) are iteratively updated. The weight matrix is uniformly initialised in [−1, 1] and scaled by 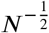, i.e. proportionally scaled by the number of units in the weight matrix.

The objective of minimising energy is captured through the following loss function:

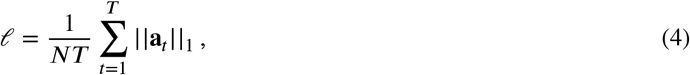

where *T* is the number of time steps, *N* is the number of units, and **a**_*t*_ is the preactivation of the units when processing the *t*-th element (or timestep) in a given sequence. We trained 10 model instances for 200 epochs on MNIST and 10 model instances for 1000 epochs on CIFAR10 with batch-size 32 (32 sequences per batch) with the Adam optimiser (*β*_1_ = 0.9, *β*_2_ = 0.999, learning rate = 0.0001) Kingma and Ba (2014). Model training differed in weight initialisation and training sequence. We tested these same 20 model instances (lesioned models in figure 4 panel A and figure 5 panel C) but with unit activity **h**_**t**−**1**_ masked such that the feedback driven units did not affect the computation of **a**_**t**_. No extensive hyperparameter search was performed. The models were trained using PyTorch 1.4.0 and Python 3.7.6.

### 4.3 Derivation and calculation of bounds on preactivation

To benchmark how well the model minimises preactivation we derived three reference measures. These were: the input activity alone (i.e. no prediction), the pixel-wise median of the entire data set removed from each image, and the pixel-wise median by image category removed from each image. To relate these benchmarks to prediction, consider that the loss function, Equation 4, measures the L1-norm of the preactivation. This preactivation is composed of the input drive and some recurrent input. Now consider that this recurrent input can be an inhibition (i.e. negatively signed) which cancels the input drive and minimises this loss. Given this consideration, we can state the three benchmarks for performance, *e*_1_, *e*_2_, *e*_3_, as (inhibitory feedback) prediction of three forms:

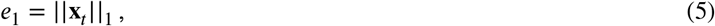

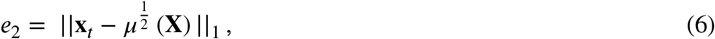

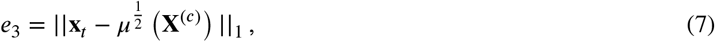

where **X** ∈ ℝ^*N*×*M*^ is the entire data set (of *M* images), 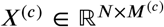 denotes the *M*^(*c*)^ data points that belong to (image) category *c* and 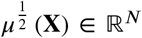 denotes the pixel-wise median of a data set **X**. In this formulation, e_1_ reflects the loss computed based upon input activity induced by **x**_*t*_ without any attempt of suppression by the RNN, *e*_2_ is the loss based upon residue activity left after subtracting the (pixel-wise) median intensity of the entire data set, and *e*_3_ is the loss based upon residue activity left after subtracting the (pixel-wise) median intensity of that image category *c*. To understand why these specific (pixel-wise median) benchmarks make sense we have to understand what the optimal behaviour is under the loss function.

Our loss function, Equation 4, is an L1 norm on the preactivation of the units of our network. This preactivation consists of a (fixed) input drive which is modified by recurrent inputs. We can therefore consider, what form of recurrent contribution would minimise this L1 norm. Lets suppose that our model (and therefore recurrent contribution) has perfect knowledge of the category of the upcoming input, labelled below *c*. Disregarding the temporal element of our RNN, and considering minimisation of a single timestep input, let us label **b** ∈ ℝ^*N*^ as our estimation of the recurrent (inhibitory) contribution. The loss, summed across all samples of the category *c*, is given by:

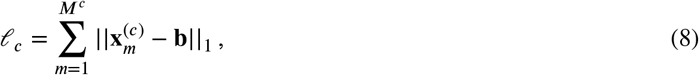

where *m* now indexes the set of examples of stimuli in category *c* (stimuli labelled 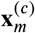) of which there are a total of *M*^(*c*)^ samples. Note that this summing is required since despite knowledge of the category *c*’s upcoming presentation, the specific sample from within this category is unknown and all samples are equally likely. Thus, to minimise the loss across all possible outcomes we minimise their summation.

The value for the recurrent contribution (**b**) that minimises this equation is given by:

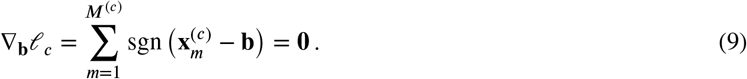

This equation is satisfied when the number of cases in which the data is greater than the recurrent contribution 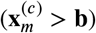 is precisely equivalent to the number of cases in which the data is smaller than the recurrent contribution 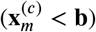. This must be true at a pixel-wise level (i.e. for each element of the vectors 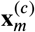 and **b**). These inequalities are consistent when our estimation is equal to the the median pixel value (per pixel) of the respective category in the data set 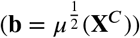. Hence, this is the prediction which would minimise our L1 loss function given the constraint of random sampling of images from each category (our benchmark *e*_3_ above). The above derivation supposed that the model had perfect knowledge of the category of upcoming samples. However, if the model was not capable of identifying which category it should expect, the best it could do would be to predict the median of the entire data set. This forms another of our benchmarks on performance (*e*_2_).

The model preactivations in Figures 2 panel A and 4 panel A are averaged over 891 sequence presentations from the test data. The benchmarks that we have produced here, *e*_1_, *e*_2_, *e*_3_, are computed over the entire training data set (60,000 MNIST Images), or per category (as in *e*_2_), and the outcome of these estimations are applied to the individual digits in each of the (891) sequences and averaged by time-step. Each data point of model preactivation represents the mean over 10 model instances. Error bars are determined with 95% confidence intervals, bootstrapped with replacement over 10000 repetitions. Each data point of the theoretical bounds is likewise determined over 10 model instances with 95% confidence intervals, bootstrapped with replacement over 10000 repetitions. The variance in the bounds is due to the variation in input sequence generation and presentation across the 10 models. The variation for model preactivation is in addition to that driven by variation in weight initialisation

The model preactivations for CIFAR10 in Figure 5 panel C are generated in the same way as for MNIST (992 sequences from CIFAR10 test set). The benchmarks *e*_1_, *e*_2_, *e*_3_, are computed over 50,000 CIFAR10 training images.

### 4.4 Determination of prediction and error units

In order to determine whether units are predictive or signal error we record their median preactivations for each category at the final time step of a sequence, when the dynamics of the network are most stable. We construct 99% confidence intervals around this median. To obtain a 99% CI we first compute the median absolute deviation around the category median preactivation. Suppose 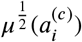 is the category median preactivation at the final time point for a particular unit *i*, then we can compute the MAD for a particular category and unit as follows:

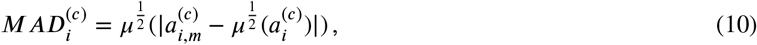

where 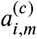 is the preactivation of unit *i* for an arbitrary sample *m* of category *c*. This means that you can find the median absolute deviation for a particular unit and category by taking the median of the absolute deviations of the unit sample preactivations from the median preactivation.

CI’s can only be analytically calculated for Gaussian distributed random variables using the standard deviation around the mean, so we have to use an approximation. We can do this by converting the obtained MADs to pseudo standard deviations (Rousseeuw and Croux, 1993) in the following way:

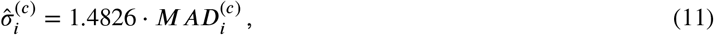

where 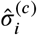 is the pseudo standard deviation around the median preactivation for unit *i* and category *c* and 1.4826 is the scaling constant for Gaussian distributed variables.

The 99% CI will then be:

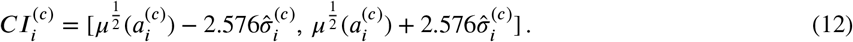

Where 2.576 is the Z-score that determines the bounds of the CI. A unit *i* will be identified as a prediction unit if 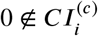 for at least one *c*.

## 5 Acknowledgements

The authors are thankful to Courtney J Spoerer for feedback and helpful discussions during an early phase of the project.

## 6 Data availability

All data and analysis code along with instructions is available at: https://github.com/KietzmannLab/EmergentPredictiveCoding

## Appendix A supplementary figures

Supplementary figures accompanying *Predictive coding emerges in energy efficient recurrent neural networks*.

**Figure A1:**
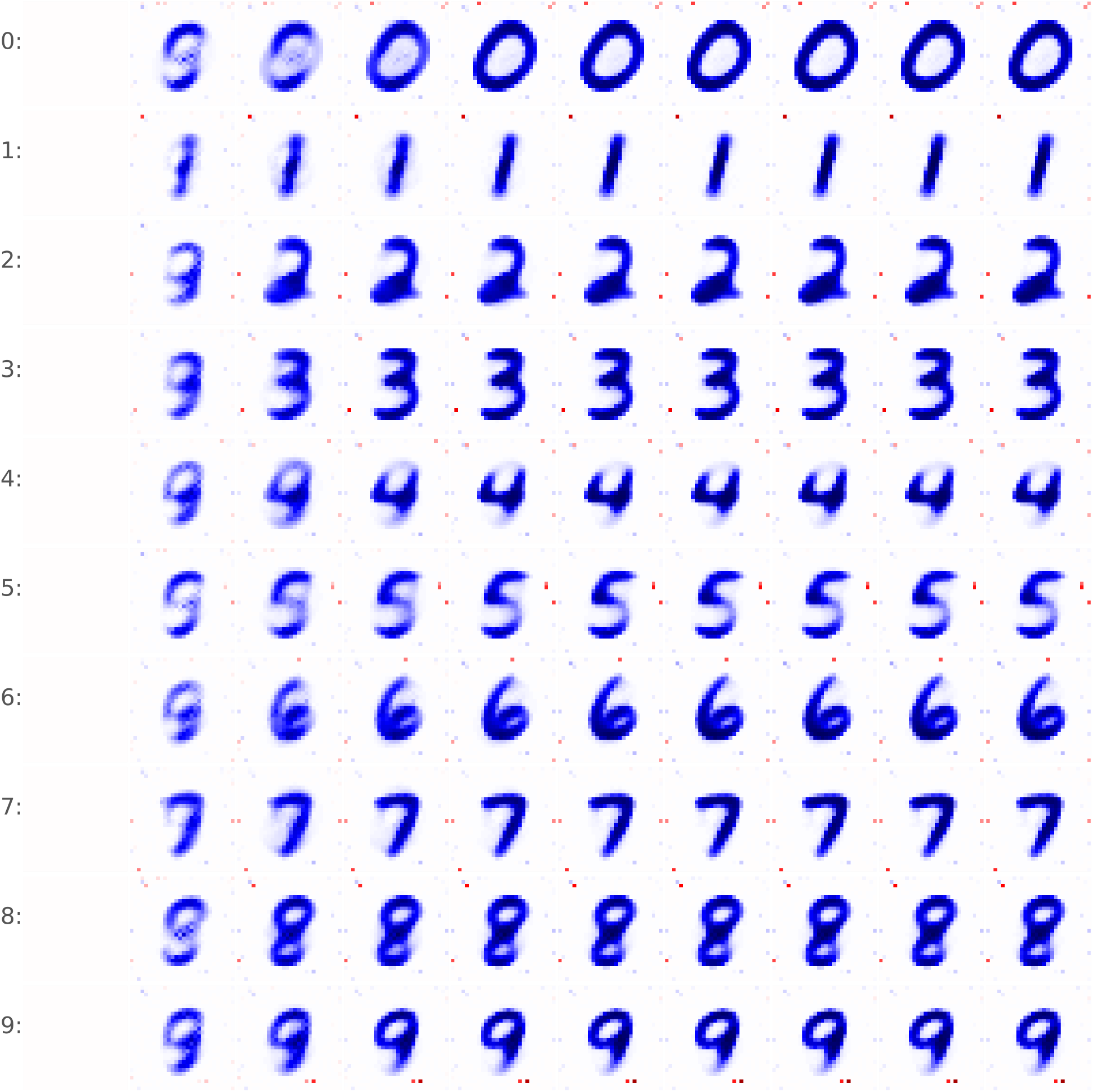
Recurrent feedback (**p**_**t**_) in the RNN for different input categories at different points in time. Each row shows the recurrent feedback at the step with a specific input category will be expected. Each column shows the recurrent feedback after a different number of preceding images in the sequence. The inhibitory effect (i.e. predictions) gets more pronounced as sequences progress.

**Figure A2:**
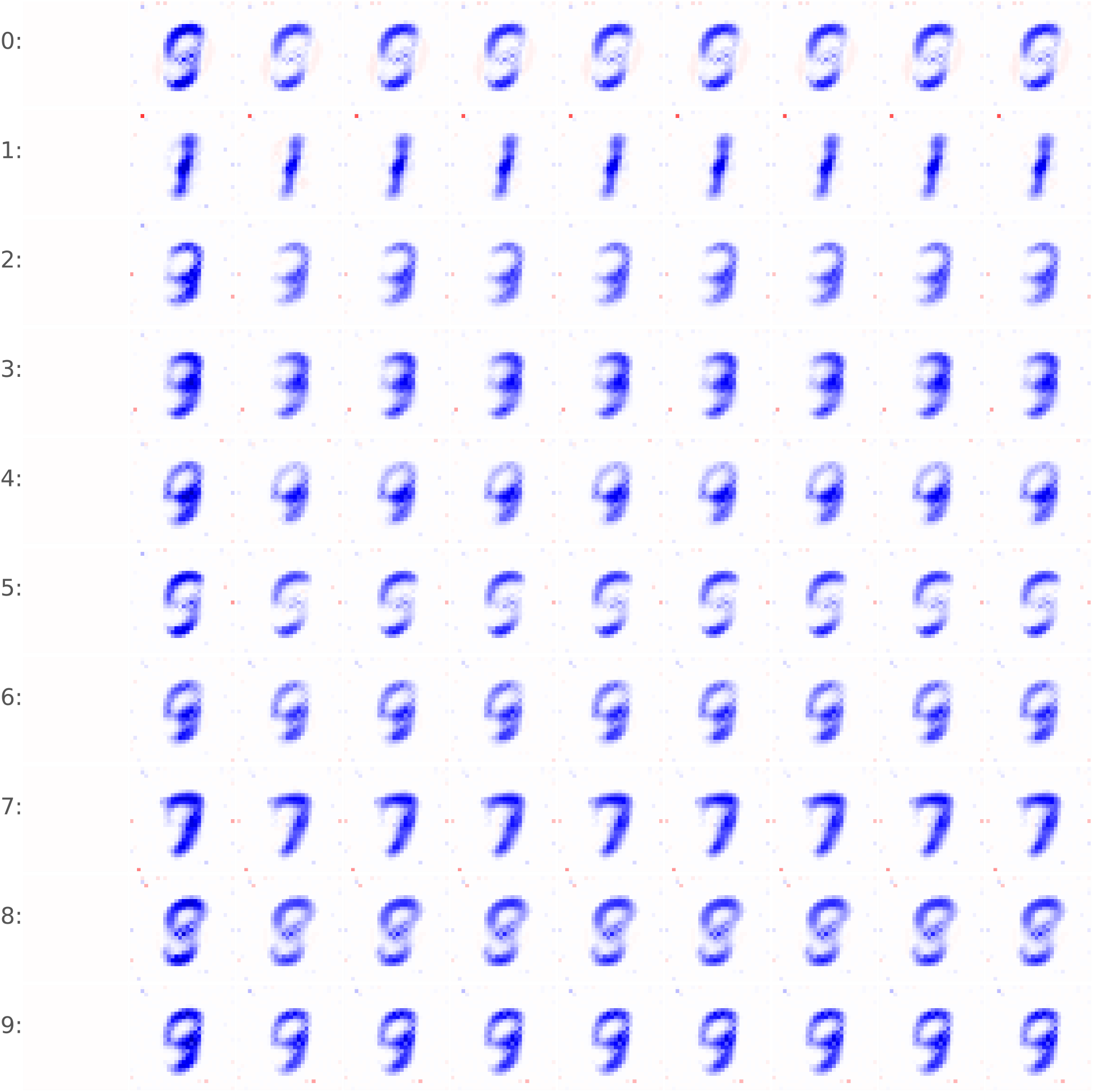
Recurrent feedback (**p**_**t**_) in the lesioned RNN model for different input categories at different points in time. Each row shows the recurrent feedback at the step with a specific input category will be expected. Each column shows the recurrent feedback after a different amount of preceding images in the sequence. The inhibitory effect with lesioned prediction units is weak and remains constant throughout the sequence.

**Figure A3:**
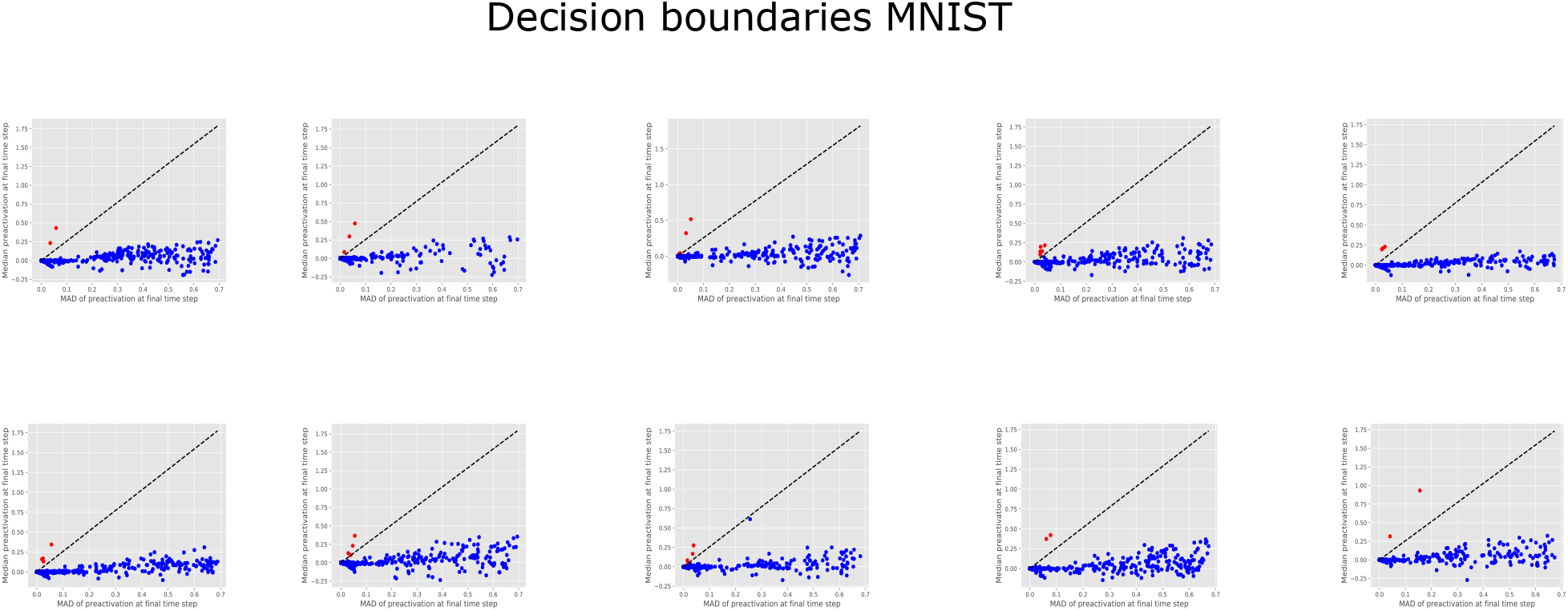
Decision boundaries for all 10 classes for the MNIST data set. Figures ordered from top left (class 0) to bottom right (class 9).

**Figure A4:**
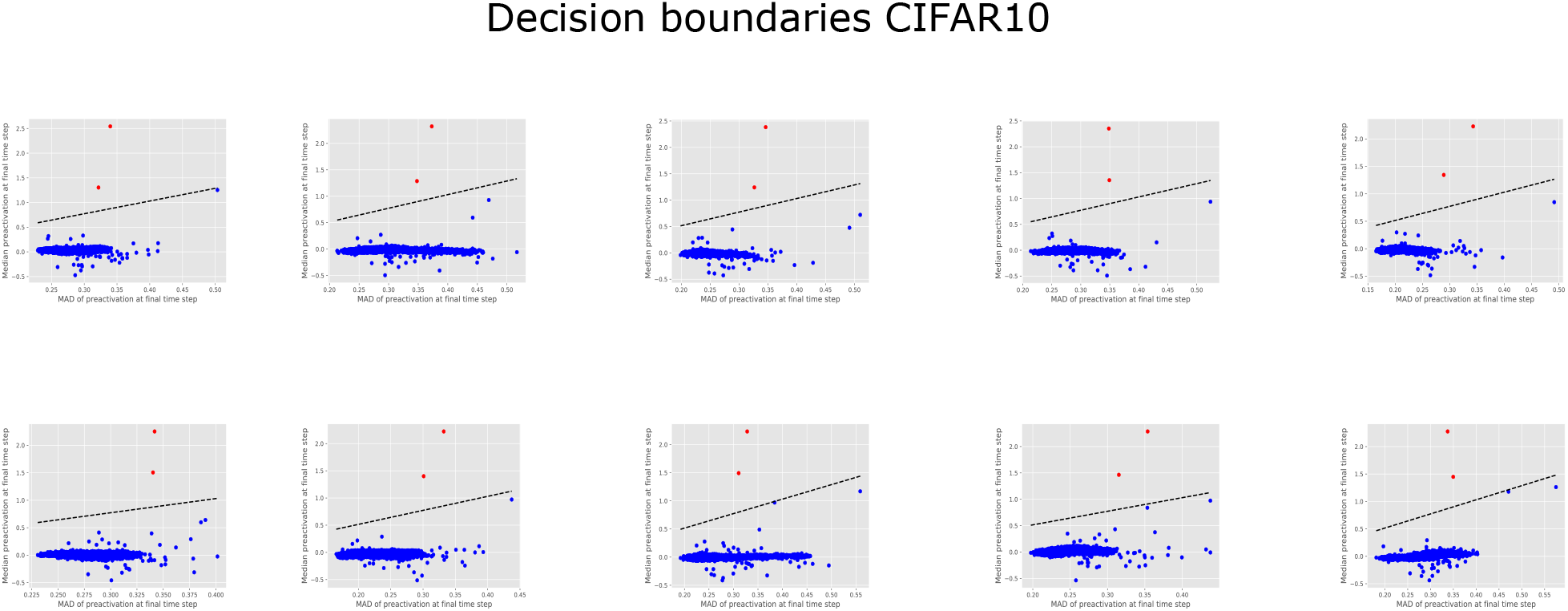
Decision boundaries for all 10 classes for the CIFAR10 data set. Figures ordered from top left (class 0) to bottom right (class 9).

## Appendix B theoretical motivation for preactivation

Supplementary text accompanying *Predictive coding emerges in energy efficient recurrent neural networks*. This supplement provides a derivation for the claim in the main manuscript that minimising preactivation leads to smaller weights and thus synaptic transmission. We do this by analysing what weight changes are induced in the case where noise causes predictions to deviate from the optimal predictions.

In this work we apply an L1 cost function to the preactivations of our units in mini batches, such that for a network of *N* units, and a given mini batch *b* consisting of *M* samples,

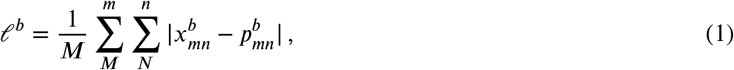

where 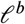 is the loss function measured for mini batch *b*. For simplicity’s sake, let us assume a batch size of one and consider a single unit within our network. This allows us to write this loss as:

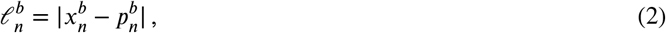

where *x*_*n*_ are the input drives to each unit, indexed *n*, and *p*_*n*_ is the recurrent network feedback which is learned in order to minimise this loss.

However, in this network we train the system to minimise the loss of prediction with some history of dynamics. Assuming that our history of dynamics are consistent but with some batch-dependent error *ϵ*, our recurrent prediction simplifies to:

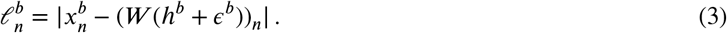

We could now plot the set of losses across all of these batches such that we would observe a distribution of losses. By carrying out SGD to minimise our loss, we would minimise the mean and the variance of this distribution.

**Figure A5:**
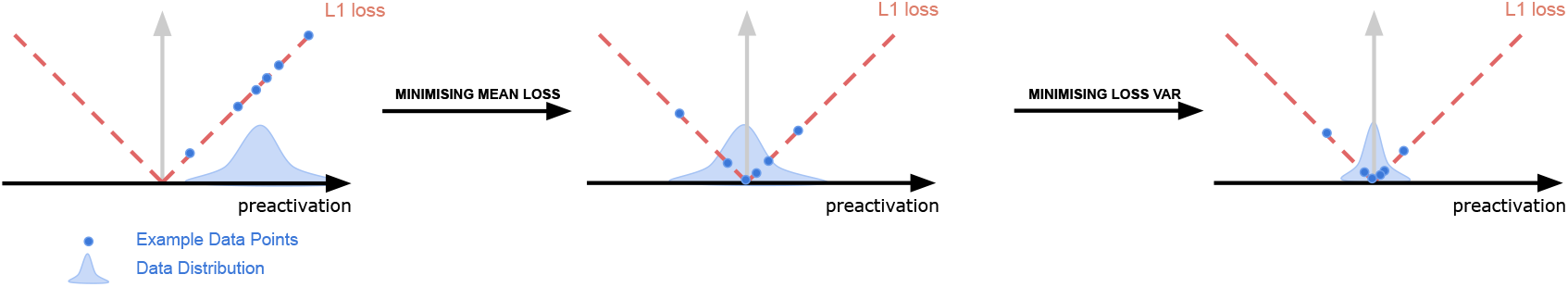
Visualisation of what happens to the loss after training. X-axis shows the preactivation for a particular data point, y-axis shows the associated loss. We can construct a distribution over the losses as well as plot the loss function to see how far the current networks performance is from optimal. Minimising the mean will not be enough to reach the optimal solution, Since the variance also contributes to the loss of network. Thus SGD will minimise the variance and the mean

## Average loss

The average loss can be computed as an expectation over samples such that

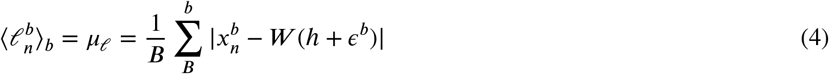

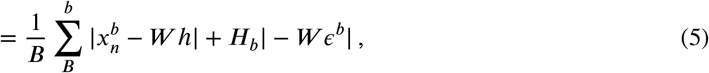

where *H*_*b*_ is an signed variable (−1/+1) depending upon whether the contribution of noise increases or decreases the magnitude of the loss, 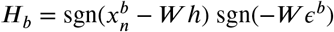 assuming no switches in signs are introduced by the noise.

Now, in computing this average, we can say that if the noise is uncorrelated (random) w.r.t. the activity and weights and has zero mean (unbiased), that it will average out such that ⟨*H*_*b*_|*W ϵ*^*b*^| = 0.0⟩ and thus

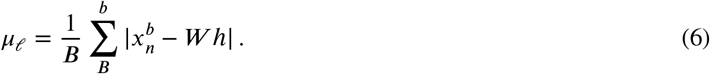

Hence, in order to minimise our average loss, we would attempt to produce prototypical predictions.

## Variance of loss

Now considering the variance of our loss

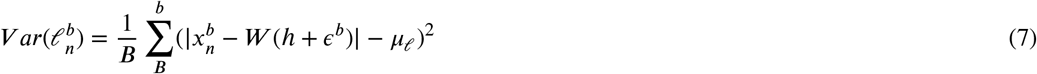

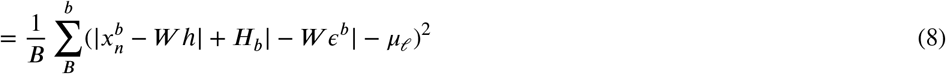

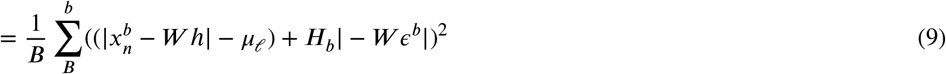

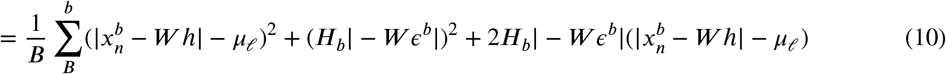

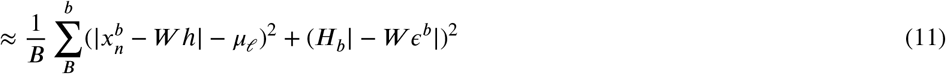

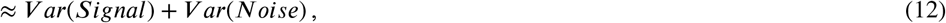

where we now have three terms (see eq 6), one term which is the variance of the noiseless predictions, a measure of the variance of noise term (scaled by W) and a cross-term correlating the noise term. Note that the correlated noise term involves multiplying two random variables, each with a mean of zero, thus we ignore this term as its expected value is zero. Finally, we are therefore left with terms of the signal and noise variance. The signal variance cannot be reduced since it is inherent to the data. The noise term however, scales with W, thus to reduce the variance caused by our noise term the weights, *W*, would be reduced. Note that the noise here refers to noise in your prediction, i.e. an uncertainty about the upcoming stimulus which is the case in real data.

